# FDTest: Fluctuation-Dissipation Theorem as a Test for Memory Effects in Brain Dynamics

**DOI:** 10.64898/2025.12.02.691886

**Authors:** Sebastian Manuel Geli, Juan Manuel Monti, Adrián Ponce-Alvarez, Gustavo Deco, Yonatan Sanz Perl

## Abstract

A central challenge in neuroscience is to understand how the brain flexibly balances local and distributed information processing to support diverse cognitive and conscious states. We hypothesize that a key signature of this balance is the presence of memory effects, which arise when a brain region’s future activity depends not only on its current state, but also on past information fed back from the wider network. Here we introduce the FDTest, a method for assessing local memory effects in multidimensional systems by measuring violations of a generalized Fluctuation–dissipation theorem (FDT). We first apply this framework to whole-brain models fitted to human neuroimaging data, showing that the brain’s memory structure reflects its underlying connectivity. We then extend the analysis to individualized models of subjects during wakefulness and deep sleep. Memory effects are consistently stronger in wakefulness, indicating richer inter-regional dependencies and more integrated dynamics. These findings establish local memory as a dynamical marker of brain state and position the FDTest as a principled tool for probing the hidden structure of neural dynamics in both models and experiments.

## INTRODUCTION

Brain function emerges from a delicate balance between localized information processing, carried out by specific regions, and distributed processing, which integrates activity across many interconnected areas [1]. This balance reorganizes dynamically to support diverse cognitive functions and conscious states. Understanding how the brain flexibly configures to produce more segregated or integrated modes of processing represents a fundamental challenge in neuroscience, with profound implications for characterizing healthy brain function [2] and identifying biomarkers of altered consciousness [3, 4] or neurological disease [5, 6]. In experimental settings, researchers have probed this balance using perturbation paradigms, such as transcranial magnetic stimulation combined with electroencephalography (EEG) recordings, where the complexity of evoked activity quantifies the interplay between functional integration and differentiation in thalamocortical networks[7–9].

We hypothesize that a key signature of this local–global balance lies in the system’s memory structure. When an element of a neural system operates relatively isolated, its future activity depends primarily on its current state, exhibiting minimal memory. Conversely, if a unit is strongly integrated into the broader network, its dynamics are shaped by information fed back through network interactions, producing local memory effects: the region’s future evolution depends not only on its present state but also on the history of interactions with the rest of the system. Perturbations make these effects visible, since a region’s response reflects the balance between its intrinsic dynamics and the hidden influence of network.

Underlying this idea is the concept of Markovianity of a stochastic process. A Markovian system is memoryless—the statistics of future states depend only on the present state and not on its past history. In principle, any closed and isolated physical system satisfies the Markov property when all microscopic variables are considered. However, a limited or coarse-grained measurement of the process may then appear non-Markovian, because the available description is insufficient to predict its future evolution [10]. At the microscopic level, neuronal firing has been modeled as a random process with a finite correlation time, thereby having a Markovian structure that can be modeled using the Fokker-Planck for-malism [11–13]. At the macroscopic level, however, the activity of brain regions exhibits long-range temporal correlations [14], reflecting the collective dynamics of distributed neuronal populations [15]. We hypothesize that macroscopic memory effects provide a window into the brain’s functional organization.

To test this, we introduce the FDTest, a method to assess local memory effects in whole-brain dynamics by detecting violations of the Fluctuation-Dissipation Theorem (FDT). Originally proposed for systems near thermodynamic equilibrium [16, 17], fluctuation-dissipation theorems were subsequently generalized for a vast class of systems, including non-Hamiltonian systems operating far from equilibrium [18, 19]. By measuring how spontaneous fluctuations deviate from the system’s response to external perturbations, FDT violations can reveal hidden memory effects that arise when the observed variables do not fully capture the underlying dynamics. Previous work has linked FDT violations to nonMarkovianity, testing this principle in one-dimensional systems [20]. Here we extend this approach to high-dimensional dynamics relevant to brain activity.

We first validate the method in a two-dimensional noisy Stuart-Landau oscillator, then apply it to a network of coupled Stuart-Landau oscillators modeling large-scale brain activity [21]. Finally, we analyze subject-specific brain models across wakefulness and deep sleep. We find that memory effects, captured by FDT deviations, are consistently stronger during wakefulness than during deep sleep and vary systematically across brain regions. These findings establish deviations from Markovianity as a quantitative measure of functional integration, where stronger memory effects reflect tighter coupling of a region to the broader network dynamics. Beyond their theoretical significance, memory effects emerge as promising biomarkers of consciousness states, with direct applications in experimental perturbation paradigms.

## RESULTS

### FDTest overview

In Fig. 1 we show a schematic representation of the methodological framework. The study consisted of two main objectives. First, we proposed and validated a framework to assess local memory effects in a high-dimensional brain model by measuring violations of the Fluctuation-Dissipation Theorem (FDT). Second, we investigated whether FDT deviations could act as a distinctive dynamical signature capable of differentiating between states of consciousness.

**FIG. 1.**
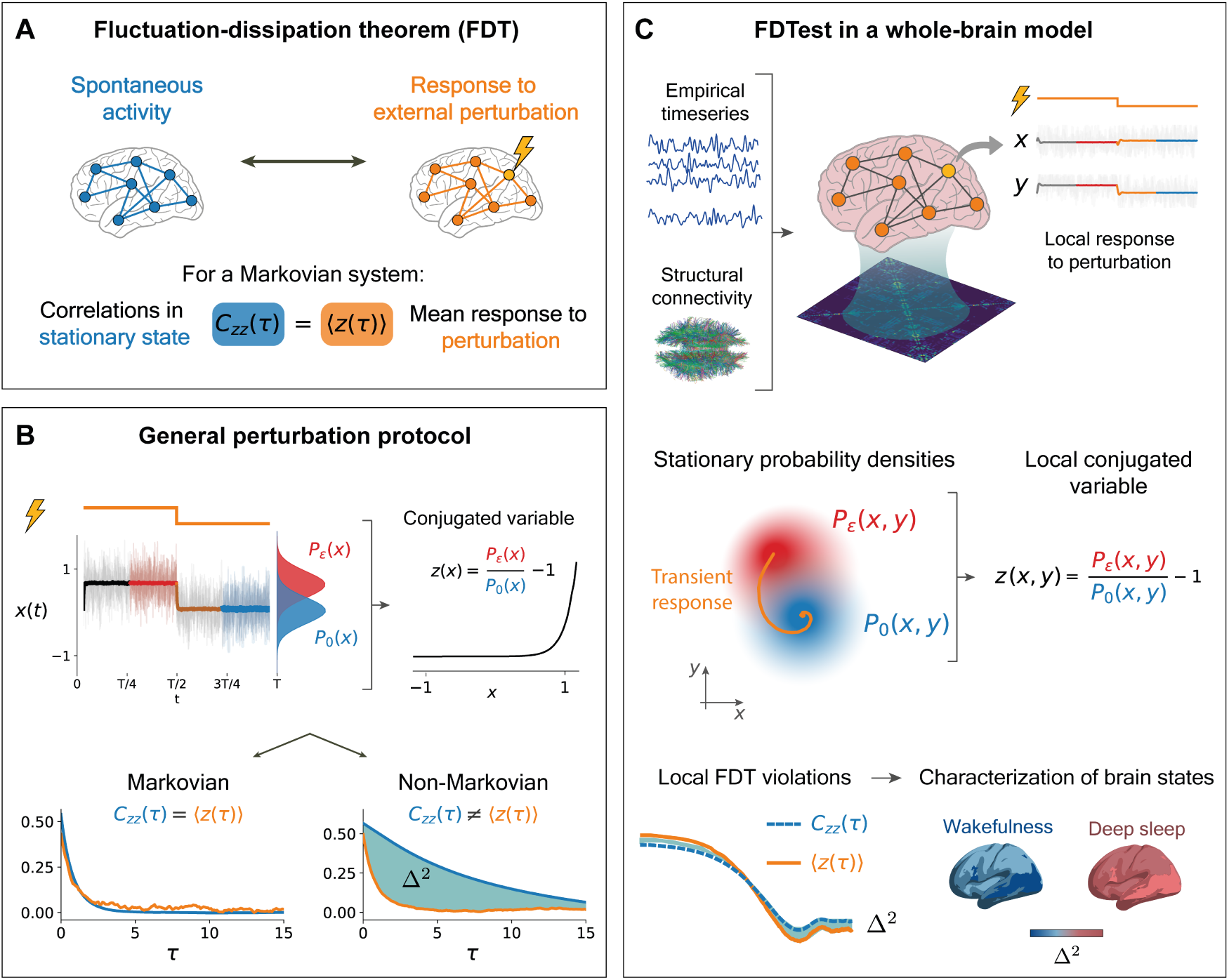
Methodological overview. **A.** The fluctuation-dissipation theorem (FDT) relates a system’s spontaneous fluctuations to its mean response to perturbations. In the formulation used here, this equality holds for memoryless (Markovian) systems, where the mean response equals the steady-state autocorrelation. **B.** Testing the FDT: a static perturbation drives the system to a perturbed steady state. After switching it off, the system transitions to a new stationary state. Stationary probability densities (P_ε_, P_0_) are estimated to define a conjugate variable *Ƶ*.The FDT holds for Markovian dynamics but is violated when relevant variables are hidden, revealing memory effects. The violation is quantified by the integrated relative difference (Δ^2^) between autocorrelation and the mean response. **C.** The FDTest assesses local memory effects in a whole-brain Hopf model constrained by anatomical connectivity and fitted to neuroimaging data. Each brain region is represented by two state variables (x, y), or equivalently by a complex variable. From these, we compute *Ƶ*(x, y), its mean transient response, and steady-state autocorrelation. Their mismatch (Δ^2^) measures regional FDT violation, mapping the brain’s memory structure across wakefulness and deep sleep (N3).

A system is said to be memoryless or Markovian when the statistics of future states depend on the present state, but not on past history. Such systems constitute an important class of stochastic models widely employed to describe systems both in and far beyond thermodynamic equilibrium. In the framework proposed here the Markovianity of a system is assessed by quantifying violations of the Fluctuation-Dissipation Theorem (FDT). In general, the FDT relates the spontaneous fluctuations of a system to its response under an applied perturbation (Fig. 1A). In the generalized relation considered here, correlations measured during spontaneous activity of a Markovian system are directly connected to the mean response elicited by an external perturbation [20]. Deviations from this relation therefore can serve as a test for the presence of memory effects (FDTest), revealing that the observed dynamics cannot be fully captured within a Markovian description.

The procedure to identify FDT violations is outlined in Fig. 1B: a system is perturbed with a static stimulus of arbitrary magnitude for a sufficient time to reach a steady state. Then, the perturbation is removed, and the system evolves to an unperturbed steady state. The probability densities are measured during the stationary states in the presence (*P_ε_*) and absence (*P*_0_) of the perturbation. Then, a new variable *Ƶ* is defined by the transformation:

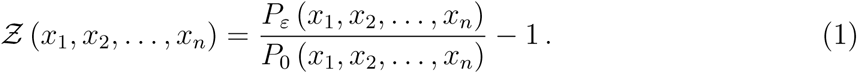

If the system is Markovian, the *Ƶ* variable obeys the FDT relation:

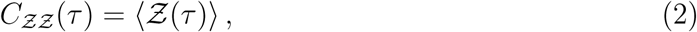

where the left-hand side of equation (2) is the autocorrelation of *Ƶ* in the absence of the perturbation, and the right-hand side is the mean transient evolution of *Ƶ* after the removal of the perturbation. The validity of this relation indicates that the system’s spontaneous fluctuations contain sufficient information to predict its mean response following an external perturbation. Conversely, deviations from this relation are indicative of non-Markovian dynamics or memory effects, where the spontaneous fluctuations of the state variables are insufficient to predict the response to a perturbation. We quantified these deviations using the relative squared difference between the two FDT terms (Δ^2^), which provides a measure of memory effects.

We first validated the FDTest in a 2-dimensional dynamical system: a noisy Stuart-Landau oscillator. To test whether the method can detect memory effects arising from missing information, we applied the FDTest under two conditions: first including all state variables in the analysis, and then systematically omitting one variable to simulate incomplete observations of the system’s state.

Then, we adapted the formalism to study a high-dimensional system: a whole-brain model consisting of a network of coupled Stuart-Landau oscillators simulating resting-state brain activity (1C). In this model, each node represents a brain region whose dynamics are described by two variables (*x, y*), with *N* = 90 regions corresponding to the AAL90 parcellation [22]. We fitted the model’s connectivity to empirical covariances matrices obtained from fMRI time series, constrained by the anatomical connectivity.

Each region’s response to external perturbations varies due to its local dynamics and network interactions. By applying the FDTest independently to each node, we mapped the memory structure across the brain: quantifying, for each region, how well its future evolution can be predicted from its spontaneous fluctuations alone, without incorporating information from other regions. This approach mimics experimental neuroscience paradigms where localized or global perturbations are applied, while the resulting activity is recorded across multiple sites [23].

Finally, we leveraged the FDTest to investigate differences in memory structure between conscious and unconscious brain states. We analyzed individualized whole-brain models for 15 subjects during wakefulness and 15 subjects during deep sleep (NREM3 stage). For each subject and condition, we applied the FDTest to calculate local FDT deviations across all brain regions. We then computed mean deviations for each brain region across subjects within each group, constructing spatial maps of memory effects, enabling the comparison of dynamical regimes between conscious and unconscious states.

### FDTest for a single stochastic Stuart-Landau oscillator

To illustrate how the FDTest can be used to detect memory effects, we first considered a two-dimensional system, a Stuart-Landau oscillator with white Gaussian noise (Fig. 2A). This is a canonical model in dynamical systems theory, and is often used in neuroscience to describe the macroscopic dynamics of an isolated brain region [21].

**FIG. 2.**
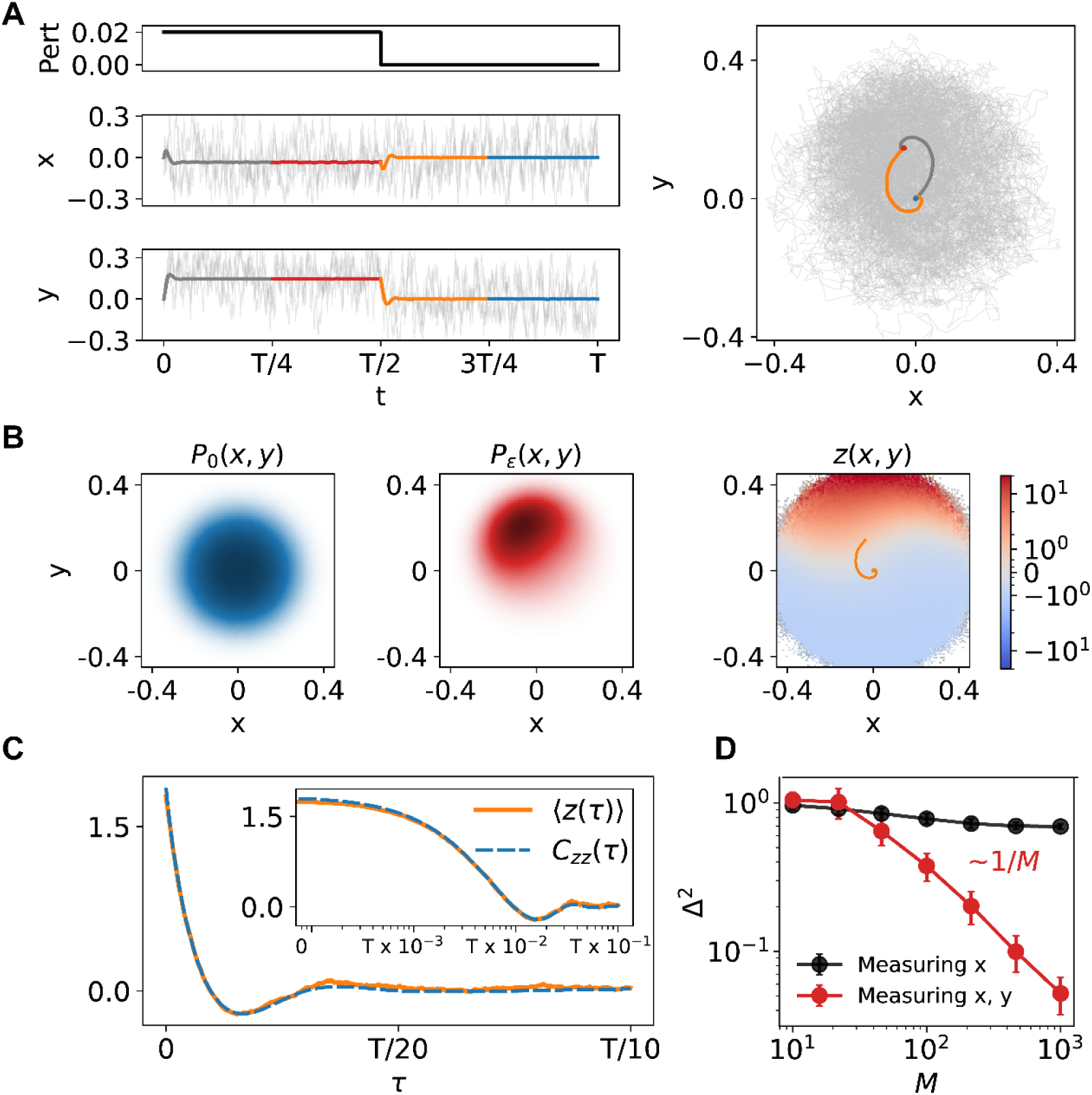
FDTest for a single stochastic Stuart-Landau oscillator. **A.** Dynamics of the perturbed oscillator. During the first half of the simulation, a constant perturbation shifts the system’s fixed point, producing a perturbed steady state (red line). At t = T/2, the perturbation is removed, and the system relaxes through a transient phase (orange) to a new steady state (blue). **B.** Stationary probability densities with the perturbation off (left) and on (center). The right panel shows the conjugated variable *Ƶ*(x, y) = P_ε_/P_0_ −1. **C.** The mean transient response to perturbation removal (orange) matches the stationary autocorrelation (blue dashed), as predicted by the FDT for Markovian systems. **D.** FDT violation Δ^2^, defined as the relative mean squared difference between ⟨*Ƶ*(τ)⟩ and C*_*ƵƵ*_*(τ), as a function of simulation count (M). With both variables measured, Δ^2^ decays as a power law; with only one, it saturates at a nonzero value, indicating persistent FDT violations and memory effects due to incomplete observation of the system.

The system was perturbed by a constant term in both equations, resulting in a shift of the system’s fixed point. We subdivided the time window *T* into four periods. The constant perturbation was active during the first two segments and then turned off at time *t* = *T/*2.

In the first and third quarters, the system undergoes a transient state in response to the change in the perturbation. In the second and fourth quarters, the system has approximately reached two different stationary states. We estimated the stationary probability densities corresponding to the states with the perturbation on (*P_ε_*(*x, y*)) and off (*P*_0_(*x, y*)) as shown in Fig. 2B, by building histograms from the ensembles of all the trajectories during the steady phases. Using the calculated densities, we estimated the conjugated variable *Ƶ*(*x, y*):

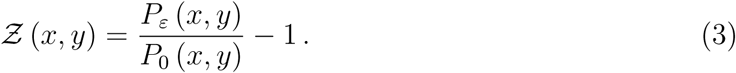

By interpolating over the region of interest in the *xy* plane, the *Ƶ* variable becomes a scalar function that assigns a real value for each (*x, y*) pair. Thus, every 2D trajectory (*x*(*t*)*, y*(*t*)) can be represented as a single time series *Ƶ*(*t*). We calculated the mean transient response to the perturbation removal by taking the ensemble average of *Ƶ*(*t*) in the third segment, which is the right-hand side of the fluctuation-dissipation relation in Eq. 2. Finally, by considering the ensemble of trajectories in the fourth segment, we can compute the stationary autocorrelation in the absence of the perturbation (left-hand side of the FDT in Eq. 2). When the number of simulations (*M*) is sufficient, these two functions overlap (Fig. 2C). We measured the discrepancy between *C_*ƵƵ*_*(*τ*) and ⟨*Ƶ*(*τ*)⟩ by computing their relative mean squared difference (Δ^2^), which quantifies FDT violations. As *M* increases, Δ^2^ decays as a power law 1*/M* (Fig. 2D, red line).

As a test case, we investigated what happens when only one variable is experimentally accessible. We replicated the previous analysis, considering only the *x* variable, omitting the *y* variable. In contrast to the complete description of the system, the FDT violation persists as the number of simulations increases. Adding more simulations does not reduce the discrepancy between *C_*ƵƵ*_*(*τ*) and ⟨*Ƶ*(*τ*)⟩ (Fig. 2D, black line). The persistent FDT violation indicates memory effects arising from a partial description of the system, as a one-dimensional projection cannot capture the full Markovian dynamics of the Stuart-Landau oscillator.

This finding highlights a key insight: FDT violations quantify the extent to which the selected variables provide a complete description of the system’s dynamics. This opens the possibility of using the FDTest to probe the system: by applying it to different subsets of variables, we can systematically map the system’s memory structure and identify hidden dependencies. In the following, we show how, in the context of whole-brain dynamics, the FDTest can characterize functional integration by assessing how local dynamics are shaped by broader network interactions.

### FDTest for coupled Stuart-Landau oscillators in a whole-brain network

We then applied the FDTest formalism for a set of coupled Stuart-Landau oscillators in a network model capturing the human whole-brain dynamics, known as the Hopf whole-brain model [21]. We treated each oscillator as a separate observable of the system and applied the procedure previously outlined. We first estimated the probability densities in the 2D space of each oscillator. Then, we computed the local conjugated variable, and calculated the mean transient response and the autocorrelation. Finally, we measured the FDT deviation by calculating the relative mean squared difference (Δ^2^) between ⟨*Ƶ*(*τ*)⟩ and *C_*ƵƵ*_*(*τ*).

We obtained heterogeneous FDT violations across the network. In particular, we found that peripheral nodes yield lower Δ^2^ values than highly connected nodes that play a central role in the network (Fig. 3A)—for example, the olfactory cortex at the network periphery exhibited a low FDT deviation, while more central and integrated regions, such as the superior frontal region, showed higher deviations. To quantify this, we examined the dependence between Δ^2^ and each node’s strength (defined as the total coupling weight) and found that more strongly connected regions systematically display greater FDT deviations (Fig. 3B). We confirmed that this effect arises from the influence of network interactions—not just from variations in the local dynamics—by comparing the results with disconnected single nodes matched for effective bifurcation parameters. The dependence could not be explained without including network dynamics (see Appendix).

**FIG. 3.**
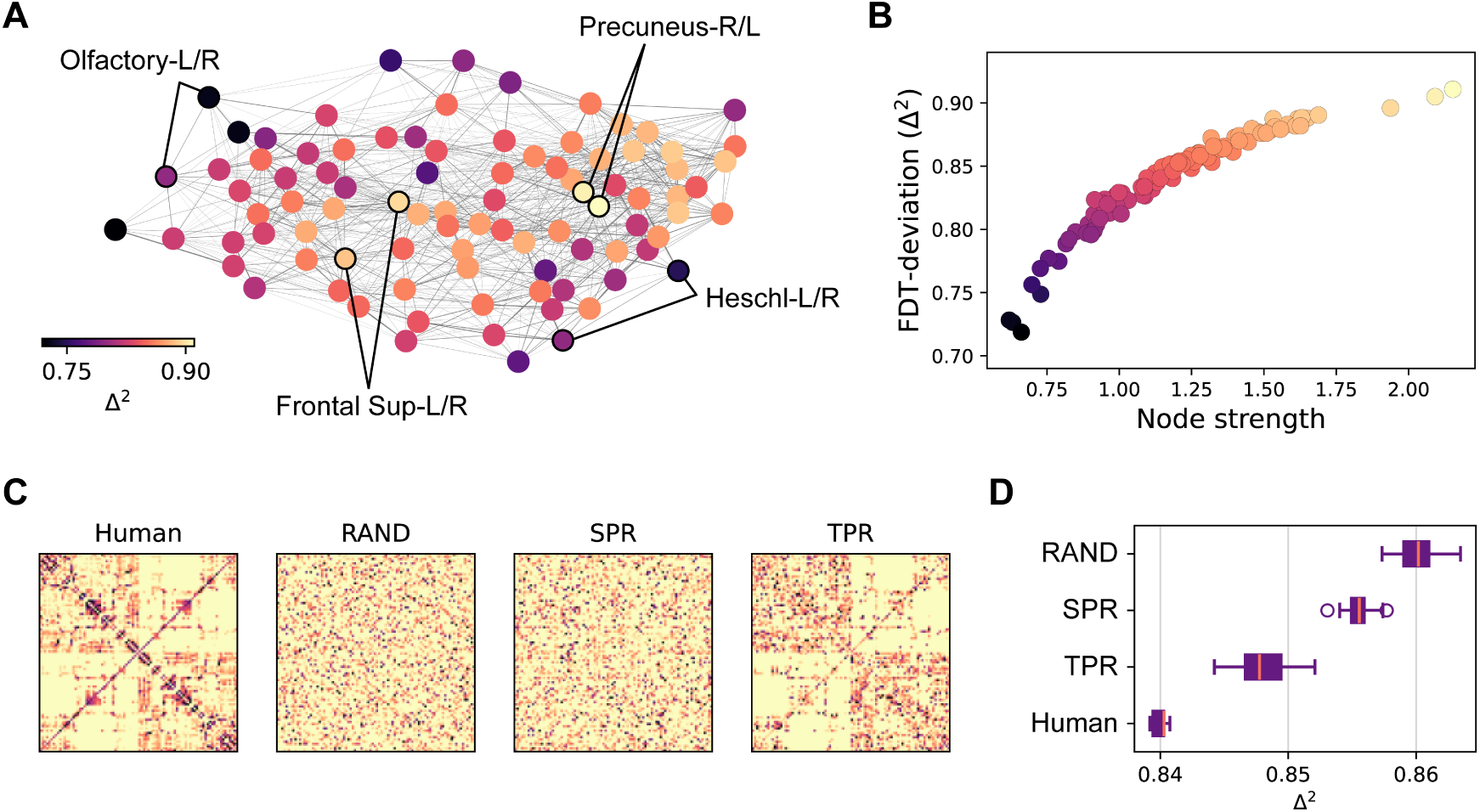
FDTest for coupled Stuart-Landau oscillators in a whole-brain network. **A.** Graph representing the fitted brain effective connectivity for an average subject, plotted using a force-directed representationt, in which edges act as springs holding nodes together and nodes repel each other, allowing visualization of the network’s center and periphery. Nodes are color-coded according to their FDT deviations, illustrating that central nodes (such as the precuneus or the superior frontal regions) exhibit higher deviations than peripheral nodes (such as olfactory regions or auditory regions like the Heschl areas). **B.** FDT deviation as a function of the node connectivity strength (total coupling weight). Nodes with higher strength show higher FDT deviations than weakly connected nodes. **C.** Null network models are obtained by rewiring the connectivity while preserving specific features (weight distribution, node strength sequence, or topology). **D.** Mean FDT deviation across the null network models, illustrating that surrogate networks are unable to reproduce the memory structure emerging from the human brain connectivity.

Having established that the FDT deviations are attributable to network properties rather than solely to local node dynamics, we asked whether these FDT deviations are distinctive of the brain network. In other words, are the FDT deviations in the brain significantly different from what would be expected by chance in networks with shared features? If that is the case, which network features are responsible for generating these specific deviations? To address this problem, we generated three sets of null network models, each preserving different features of the original brain connectivity: weight distribution (RAND), node strength sequence (SPR), and topology (TPR). For each network, the global FDT deviation was calculated as the average Δ^2^ across all nodes.

The key finding from this analysis is that surrogate networks are unable to reproduce the distributions of FDT violations observed in the human brain connectivity (Fig. 3D).

Specifically, the RAND null model, which was obtained by unconstrained rewiring of the original network, represented the least conservative model and yielded the highest FDT deviation values.

The SPR null model was constructed through random rewiring, while preserving the strength of each node, which we previously found to be closely related to FDT deviations. These networks yielded FDT deviations closer to those of the original connectivity, but still significantly different, showing that node strength alone is insufficient to explain the memory structure of the human brain network.

Finally, the TPR model was obtained by a random rewiring constrained by the network topology: the weights were permuted without moving the edges. This was the set of networks that produced FDT deviations closest to the original network, although they remained significantly higher.

### FDTest for subject-specific models of different brain states

We fitted whole-brain models to neuroimaging data from human participants during different brain states: wakefulness and deep sleep (N3). We then applied the FDTest to each individualized model and obtained the FDT deviations across all brain regions for each subject.

In Fig. 4A, we show the distributions of the average deviation across brain regions for all subjects during the two conditions. Subjects exhibited significantly higher FDT deviations during wakefulness than during deep sleep. Considering the full cohorts of 15 participants each, the Wilcoxon rank-sum test yielded *p <* 0.008. Additionally, for the subset of 12 participants that completed both sessions, we performed a paired test (Wilcoxon signed-rank test), which yielded *p <* 0.02.

**FIG. 4.**
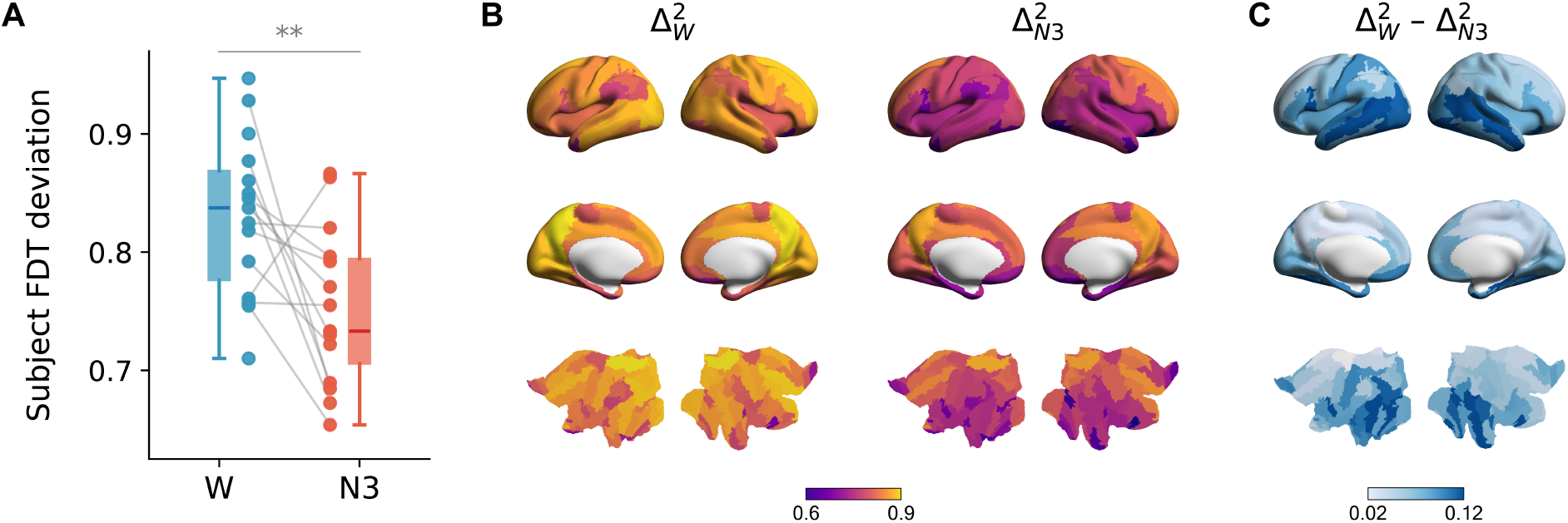
FDTest for subject-specific models of different brain states. **A.** Mean FDT deviations across brain regions for all subjects during wakefulness (W) and deep sleep (N3). Local FDT deviations are significantly higher during wakefulness, indicating stronger memory effects. **B.** Topographic map of memory effects averaged across subjects for each condition. Brain regions are colored according to their Δ^2^ values, measuring the magnitude of local FDT violations. **C.** Difference in local memory effects between wakefulness and deep sleep 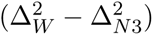. Darker blue regions indicate larger FDT violations during wakefulness compared to N3 sleep, with the largest changes observed in temporal and occipital areas.

In Fig. 4B we show brain maps of local memory effects during wakefulness and deep sleep, obtained by averaging the local deviations across subjects. In Fig. 4C, we display the difference between wakefulness and deep sleep deviations. Our results indicate that, across brain regions, FDT deviations are higher during wakefulness. The largest changes are observed in temporal and occipital areas.

Finally, we explored the dynamic correlates of FDT deviations. First, we studied the stability of each fitted model (Fig. 5A). For each subject, we linearized the model around the fixed point and calculated the eigenvalues of the Jacobian matrix. All models had only eigenvalues with negative real parts, indicating stability. We compared the spectral abscissa, which is the greatest real part of the set of eigenvalues. The closer this value is to zero, the slower the system decays to equilibrium. We found a positive correlation between the spectral abscissa and the mean FDT deviation across nodes in the system (Pearson *r_W_* = 0.98, *p <* 3 × 10^−10^ for wakefulness and *r_N_*_3_ = 0.79, *p <* 5 × 10^−4^ for deep sleep), indicating that system instability is associated with higher deviations. However, we found no significant difference in the stability between wakefulness and deep sleep (Fig. 5B), indicating that this factor does not explain the difference in FDT deviations between conditions (*p* = 0.4).

**FIG. 5.**
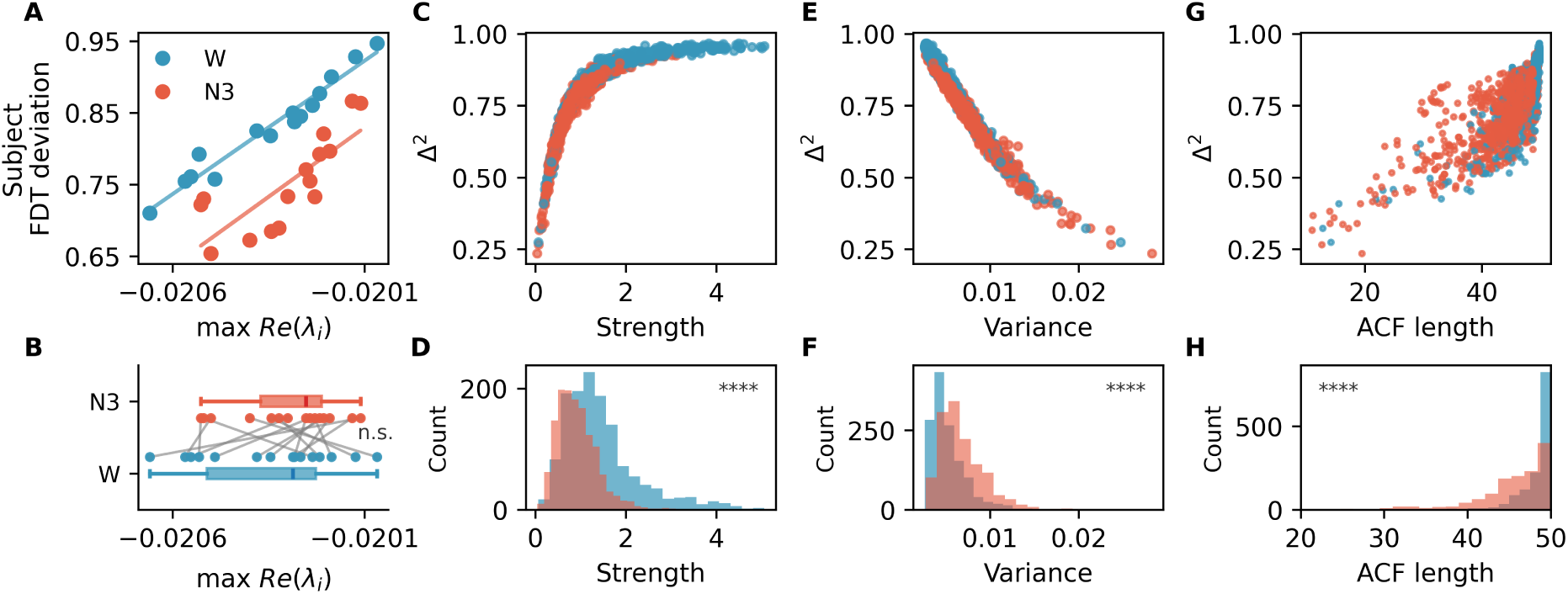
Dynamic correlates of FDT deviations. **A.** Subject-level FDT deviations are positively correlated with the real part of the eigenvalue closest to zero from the linearized Jacobian (*r_W_* = 0.98*, p <* 3 × 10^−10^ for wakefulness and *r_N_*_3_ = 0.79*, p <* 5 × 10^−4^ for deep sleep). Systems whose fixed point is closer to losing stability show larger FDT deviations. **B.** Real parts of the eigenvalue closest to zero for all fitted models, showing no significant difference in stability between wakefulness and deep sleep. **C.** Regional FDT deviations increase with the node’s effective connectivity strength. **D.** Connectivity strengths are significantly higher during wakefulness than during deep sleep. **E.** Regional FDT deviations are higher for nodes with smaller variance. **F.** Node variance is significantly higher during deep sleep than during wakefulness. **G.** Regional FDT deviations are higher for nodes with higher autocorrelation lengths. **H.** During deep sleep, temporal autocorrelations decay faster than in the wakefulness state

We then examined the dependence of Δ^2^ on each node’s effective connectivity strength (Fig. 5C). We found that nodes with higher connectivity strength show higher FDT deviations. This finding aligns with our previous analysis of the average resting state brain. Additionally, we observed that the strength of the learned connectivity was significantly reduced during deep sleep (Fig. 5D) (*p <* 10^−4^). Individual brain regions in the sleeping brain present weaker functional connections and thus their dynamics are better predicted by their own stationary fluctuations, yielding smaller FDT deviations.

Next, we simulated all models in the absence of perturbations and computed the variance of each node’s time series. We found that FDT deviations decrease as a function of node variance (Fig. 5E). Indeed, nodes with higher connection strength exhibited lower variance. Previous human and animal studies have also shown experimentally that high signal fluctuations are associated with lower functional connections [24]. Moreover, it has been demonstrated that this structure–function pattern is reproduced by the Hopf model [25]. Here, we show that such variations of the signal fluctuations create a hierarchy in FDT deviations across the network. Furthermore, during deep sleep, the variance increased significantly relative to the wakefulness state (*p <* 10^−4^; Fig. 5F), consistent with experimental observations [26–28]. Finally, we computed the autocorrelation function (ACF) of each node for all subjects. For the Hopf whole-brain model in the studied regime, the ACF envelope is approximately a decaying exponential function *ACF* (*τ*) ∼ *e*^−^*^τ/ξ^* [25]. We characterized the decay speed by fitting the autocorrelation length *ξ* (see Methods). We found that FDT deviations are larger for nodes with slower autocorrelation decays (Fig. 5G). This result aligns with previous experimental observations showing that more strongly connected nodes exhibit slower dynamics [29, 30]. It has also been shown that the variance and temporal scales of fMRI fluctuations have opposite relationships with the structural connectivity: while more strongly connected regions display slower timescales, their activity presents lower variance [25]. Here, we show an analogous pattern: larger FDT deviations occur in regions with longer autocorrelation timescales but lower variances. Lastly, we found that during deep sleep, there is a significant decrease in the temporal autocorrelation length (*p <* 10^−4^; Fig. 5H), consistent with previous experimental observations [28].

## DISCUSSION

### Memory effects as a window into the local–global balance of brain dynamics

In this work, we introduced the FDTest, a method based on the fluctuation-dissipation theorem (FDT) to probe memory effects in whole-brain dynamics. We hypothesized that memory effects in regional activity reflect the balance between local and distributed processing in the brain.

Previous studies established theoretical connections between FDT and the Markov property [20, 31]. Here we extend these ideas to high-dimensional dynamics and show that violations of FDT provide a systematic way to study the balance between autonomous dynamics and global integration in the brain. We show that memory effects offer a dynamical signature of this balance: regions that operate more independently behave close to a Markovian regime, whereas regions more strongly influenced by the network exhibit deviations from Markovianity.

Our validation in a noisy Stuart–Landau oscillator illustrates this principle: when all state variables are measured, the FDT holds, consistent with a Markovian description. However, omitting one variable leads to a violation of the fluctuation-dissipation relation, emphasizing that an incomplete set of variables fails to capture the full system dynamics and gives rise to memory effects.

Extending this logic to large-scale networks, we applied the FDTest to individual nodes in a brain model, representing distinct brain regions, thereby constructing a map of memory effects in the brain network. We found that deviations from the FDT are significantly influenced by network topology. Peripheral nodes, weakly coupled to the network, exhibit minimal FDT deviations, whereas highly connected nodes display large violations. This finding indicates that strongly connected hubs, which integrate information from multiple regions, exhibit more pronounced memory effects: their future evolution cannot be predicted solely by their own stationary dynamics; rather, it is determined by the broader network state. This aligns with previous observations that these regions display slower intrinsic timescales and lower variance, reflecting their role as integrative cores [25]. Conversely, lower-order sensory areas, optimized for encoding rapid transient responses to the external stimuli [25, 32], remain more autonomous. Our work extends these insights by showing that FDT deviations provide a dynamical marker of this hierarchy: stronger network integration is accompanied by enhanced local memory effects.

Previous studies have applied fluctuation-dissipation relations to quantify non-equilibrium dynamics in the brain [33–35]. However, those formulations rely on the assumption of thermodynamic equilibrium, so that violations of the FDT are interpreted as signatures of deviations from equilibrium. In contrast, the present formulation does not require equilibrium, but instead assumes Markovianity. Consequently, deviations from the FDT here reflect non-Markovianity or memory effects, rather than non-equilibrium behavior. This distinction is crucial, as it allows to disentangle the effects of irreversibility from those of hidden dependencies, enabling the study of memory in systems both near and far from thermodynamic equilibrium.

Finally, we remark that the FDTest can be applied directly to time series data to assess whether a given set of observables is sufficient for constructing a complete dynamical description that allows to predict the system’s evolution– a central question in data-driven models of complex systems. In neuroscience, limited observability is a fundamental constraint: most experimental setups only record the activity in a subset of regions (e.g., EEG, local field potentials). Yet modeling approaches often assume Markovian dynamics [36, 37]. The FDTest offers a principled way to assess the validity of this assumption, and it can guide the design of perturbation protocols and the selection of observables.

### Memory effects as a biomarker of consciousness

Our application of the FDTest to subject-specific whole-brain models revealed that FDT deviations are higher during wakefulness compared to deep sleep (N3). This supports our hypothesis that the brain’s memory structure reflects its functional organization: in wake-fulness, responses to perturbations cannot be predicted from the local steady state alone because regional dynamics are less autonomous and more strongly shaped by network inter-actions. By contrast, during deep sleep, the activity is more locally determined and closer to a Markovian description. These findings align with the existing literature indicating that the sleeping brain dynamics exhibit reduced dimensionality [38, 39] and complexity [40–43], as well as decreased global synchronization [44]. Our contribution adds a complementary perspective: the sleeping brain exhibits weaker memory effects because local dynamics are less constrained by network interactions, while the enhanced memory effects of wakefulness reveal richer inter-regional dependencies.

These results open new directions for the characterization of brain states. In various clinical and experimental protocols, cortical neurons are perturbed with transcranial magnetic stimulation (TMS) while the response is recorded with EEG [45, 46]. These setups are used to assess consciousness: the TMS evokes a complex pattern of activation in conscious states while reacting with a simple, localized response if consciousness is reduced [7, 8]. An established measure of such complexity is the Perturbational complexity index (PCI) [9]. Our work presents a new way of analyzing the evoked responses in such paradigms by explicitly testing for local memory effects, offering a theoretical lens on brain responsiveness and integration. The FDTest can be applied directly to data from stimulation experiments in a model-free manner, even in contexts where only a few observables are available (for example, a handful of electrodes in certain brain regions) and where global markers (e.g., functional connectivity) may be unreliable or unavailable.

Lastly, we would like to acknowledge certain limitations of our study. First, here we focused on applying the FDTest to individual brain regions. In practice, one might want to assess whether larger sets of variables (e.g., multichannel EEG recordings or a group of electrodes) jointly exhibit Markovian dynamics. Evaluating the FDTest in larger observable sets is a natural next step. However, this can present a significant experimental and computational challenge, since the required perturbations scale rapidly with state-space dimensionality. Furthermore, we validated our method using synthetic perturbations in stochastic models, where the number of simulations can be controlled. In experimental stimulation setups, typically around tens or a few hundred perturbations are performed, which constrains statistical power. Model-based perturbations can serve as an initial step to guide experimental parameters, such as perturbation intensity and the number of simulations required. Finally, future work should aim to explore different types of perturbation, other than step-like, in order to make the method suitable for various experimental setups.

## METHODS

### FDTest general protocol

For all the systems considered in this work, we followed the same perturbation protocol for quantifying FDT deviations [20]. Briefly, a system is subject to a static external perturbation that is turned off. The process of duration *T* is divided in four stages. In Stage I, the perturbation is turned on and the system evolves towards a perturbed steady state. In Stage II, the system has reached a perturbed steady state characterized by the perturbed stationary distribution *P_ε_*. In Stage III, the perturbation is switched off and the system undergoes a transient regime. Finally, in Stage IV, the system has reached a new steady state, characterized by the unperturbed stationary distribution *P*_0_. The process is repeated multiple times to have an estimate of the steady state densities (*P_ε_*, *P*_0_) and sufficient realizations of the transient evolution.

From the steady state distributions, the conjugated variable *Ƶ* is calculated:

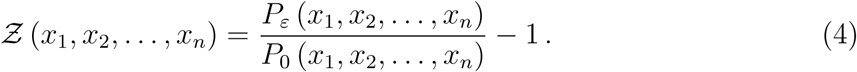

Even if the state variables define a multidimensional space, the conjugated variable is unidimensional.

From Stage III the mean transient response, ⟨*Ƶ*(*τ*)⟩ is calculated by ensemble averaging the trajectories in the *Ƶ* space. Then, from Stage IV the autocorrelation function of *Ƶ* is calculated: *C_*ƵƵ*_*(*τ*) = ⟨*Ƶ*(*t*)*Ƶ*(*t* + *τ*)⟩.

If the system is Markovian, the *Ƶ* variable obeys the FDT relation:

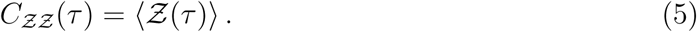

Then, the FDT deviation can be quantified as the relative squared difference between the two terms:

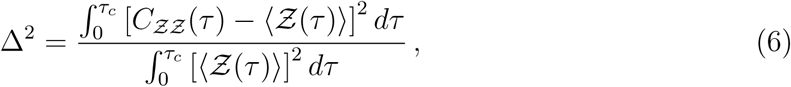

where *τ_c_*is a suitably chosen cutoff time capturing the relevant dynamical timescales of the autocorrelation and the transient regime.

### FDTest in a noisy Stuart-Landau oscillator

We first considered a noisy Stuart-Landau oscillator with bifurcation parameter *a*, frequency *ω*, and noise amplitude *σ*:

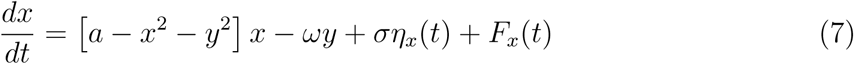

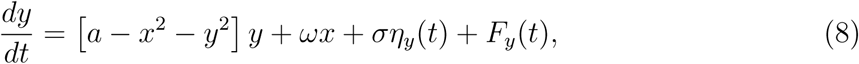

where *η_x_*(*t*) and *η_y_*(*t*) are *z*ero-mean white Gaussian noises: ⟨*η*(*t*)⟩ = 0 and ⟨*η*(*t*)*η*(*t*^′^)⟩ = *δ*(*t* − *t*^′^).

This system undergoes a Hopf bifurcation at *a* = 0. For *a <* 0 there is a stable fixed point, located at (*x, y*) = (0, 0) in the absence of perturbation. In this regime, the noise induces oscillations of the system. For *a >* 0 there exists a stable limit cycle oscillation with radius *r* = √*a* and frequency *f* = *ω/*2*π*.

The perturbation is an additive constant that is switched off halfway through the simulation in a step-like manner:

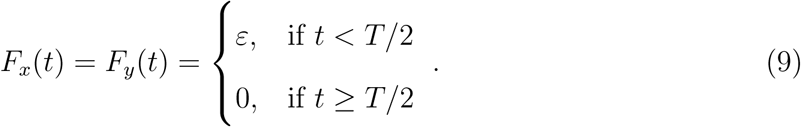

The perturbation shifts the stable fixed point of the system from (0, 0) during the first half of the simulation. For the example shown in Fig. 2, the parameters were: *a* = −0.02, *ω* = *π/*20, *σ* = 0.06, *ε* = 0.02 and *T* = 1000. The stochastic system was integrated using the Euler-Maruyama method with an integration step of *dt* = 0.1.

The steady state densities with the perturbation on and off, *P_ε_*(*x, y*) and *P*_0_(*x, y*), were estimated by constructing 2D histograms of the stationary trajectories with the perturbation on and off. Here we considered a grid of 100 x 100 bins, covering the range between the minimum and the maximum observed value in each direction. Using finer grids produces a better description, provided that the number of simulations is also increased accordingly.

The conjugated variable was evaluated in each bin following Eq. 4, and then bilinearly interpolated to calculate the mean transient response ⟨*z* [*x*(*τ*)*, y*(*τ*)]⟩ and the autocorrelation *C_*ƵƵ*_* = ⟨*Ƶ* [*x*(*t*)*, y*(*t*)] *Ƶ* [*x*(*t* + *τ*)*, y*(*t* + *τ*)]⟩. The FDT violation was quantified using a discrete form of Eq. 10, with *τ_c_* = 100:

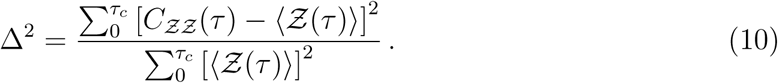

We replicated the previous procedure but considering only the *x* variable, neglecting the *y* variable. We built a 1D histogram for estimating *P_ε_*(*x*) and *P*_0_(*x*) and evaluated the conjugated variable *z*(*x*) = *P_ε_*(*x*)*/P*_0_(*x*) − 1 in each bin. Then, we did a linear interpolation and calculated the mean transient response and autocorrelation, and their normalized squared distance. We repeated the process multiple times, considering both the full set and the reduced set of variables, while varying the total number of simulations, *M*.

### Datasets

#### Structural data

The whole-brain model was initialized using structural connectivity (SC) in the AAL parcellation of 90 cortical regions [22]. This dataset was obtained from the Human Connectome Project (HCP)’s public data release in March 2017. It was acquired using a customised 3 Tesla Siemens Connectom Skyra scanner with a standard Siemens 32-channel RF receive head coil. For each participant, at least one 3D T1w MPRAGE image and one 3D T2w PACE image were collected at a resolution of 0.7 mm³. Details on the acquisition parameters are available on the HCP website [47].

Multi-shell diffusion-weighted imaging data from 32 HCP database participants (scanned for approximately 89 minutes) was used to estimate connectivity, as described by Horn et al. [48]. Briefly, the data were processed using a generalised q-sampling imaging algorithm implemented in DSI Studio (http://dsi-studio.labsolver.org/). White-matter masks were generated by segmenting the T2-weighted anatomical images, which were then co-registered to the b0 diffusion image using SPM12. Within the white-matter mask, 200000 fibres were sampled in each HCP participant. These fibres were then transformed into MNI space using Lead-DBS [49].

#### Human sleep data

##### Ethics

Written informed consent was obtained, and the study was approved by the ethics committee of the Faculty of Medicine at the Goethe University of Frankfurt, Germany.

##### Participants

We used fMRI and polysomnography (PSG) recordings from a larger database that included all four PSG stages. In this study, we focused on wakefulness and deep sleep (N3). Exclusion criteria were based on the quality of the simultaneous acquisition of EEG, EMG, fMRI, and physiological recordings. This resulted in 15 recordings for each condition, comprising data from 12 subjects who completed both conditions, along with two additional recordings per condition from subjects who provided only one high-quality session.

##### Acquisition and pre-processing of fMRI and polysomnography data

Neuroimaging fMRI data were acquired using a 3T Siemens Trio system (Erlangen, Germany). A total of 1,505 volumes of T2*-weighted echo-planar images (EPI) were collected with a repetition time (TR) of 2.08 seconds and an echo time (TE) of 30 ms. Imaging parameters included a matrix size of 64 × 64, voxel dimensions of 3 × 3 × 2 mm³, a 50% distance factor, and a field of view (FOV) of 192 mm². Preprocessing of the EPI data was performed using SPM8 (http://www.fil.ion.ucl.ac.uk/spm/), involving realignment, normalization to MNI space, and spatial smoothing with an 8 mm³ FWHM Gaussian kernel. The data were then spatially downsampled to a resolution of 4 × 4 × 4 mm³. Physiological noise components related to cardiac and respiratory activity were estimated using the RETROICOR method [50], based on simultaneously recorded ECG and respiration signals. These components, along with motion parameters, were regressed out.

The data were temporally band-pass filtered in the range 0.008-0.08 Hz using a sixth-order Butterworth filter, and timeseries were extracted in the AAL90 parcellation [22].

Simultaneous PSG was performed through the recording of EEG, EMG, ECG, EOG, pulse oximetry, and respiration. EEG was recorded using a cap (modified BrainCapMR, Easycap, Herrsching, Germany) with 30 channels, of which the FCz electrode was used as reference. Data were sampled at 5 kHz and low-pass filtered at 250 Hz.

MRI and pulse artefact correction were applied based on the average artefact subtraction method [51] in Vision Analyzer2 (Brain Products, Germany). EMG was recorded from chin and tibial sites, while ECG and EOG were recorded bipolarly, also at a sampling rate of 5 kHz and low-pass filtered at 1 kHz. Pulse oximetry was obtained via the Trio scanner, and respiration was measured using MR-compatible equipment (BrainAmp MR+ and BrainAmp ExG; Brain Products, Gilching, Germany).

Participants were instructed to lie still with their eyes closed and relax during scanning. Sleep staging was performed by a certified sleep expert based on EEG recordings, following the AASM 2007 criteria. Previous analyses using the same dataset and preprocessing pipeline have been reported [52]. The time duration of fMRI signal in each condition is dependent on the time that each participant spent in each sleep stage. Here, we considered the wakefulness and deep sleep stages, where the time duration in each condition varies from 110 to 720 volumes and from 88 to 1002 volumes, respectively.

##### Hopf model of whole-brain dynamics

We considered a Hopf whole-brain model consisting of a network of coupled Stuart-Landau oscillators. In this model each oscillator represents a brain region, and the oscillators are coupled by a generative effective connectivity matrix 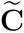 which is obtained by a fitting constrained by the anatomical connections between regions. Thus, the whole-brain dynamics are defined by the following set of coupled equations:

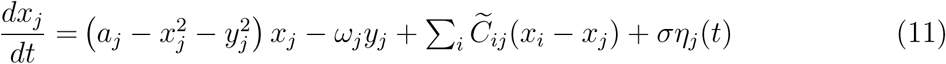

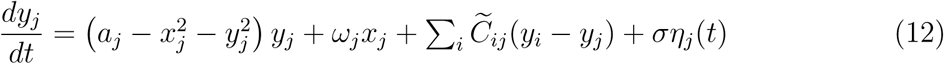

We couple the equations using the common diffusive coupling, which approximates the simplest (linear) part of a general coupling function. This approximation is valid in the weakly coupled oscillator limit, in which the coupling preserves the periodic orbit of the uncoupled oscillators.

The intrinsic frequencies *ω_j_* were estimated from the data as the subject-averaged peak frequencies of the narrowband blood-oxygen-level-dependent (BOLD) signal of each region. These frequencies lay within the 0.008–0.08 Hz range. For *a_j_ >* 0, the local dynamics exhibit a stable limit cycle, resulting in noisy, self-sustained oscillations with frequency *ω_j_/*(2*π*). For *a_j_ <* 0, the local dynamics present a stable spiral fixed-point, producing noisy oscillations.

The standard deviation of the noise was set to be the same for all nodes of the network, *σ* = 0.09. This is the averaged value across subjects and conditions found in earlier work [34].

To fit the coupling matrix 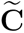, we use a pseudo-gradient descent procedure [53, 54]. Specifically, 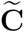 was fitted to ensure that the model accurately reproduces the empirically measured covariances **FC**^empirical^ (i.e., the normalized covariance matrix of the functional neuroimaging data) and the empirical normalized time-shifted covariances **FS**^empirical^(*τ*), where *τ* denotes the time lag. The value of *τ* was selected as the time lag that led to a decrease in the average autocorrelation. These normalized time-shifted covariance matrices are generated by taking the shifted covariance matrix **KS**^empirical^(*τ*) and dividing each pair (*i, j*) by 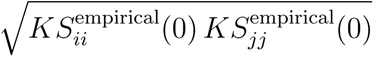

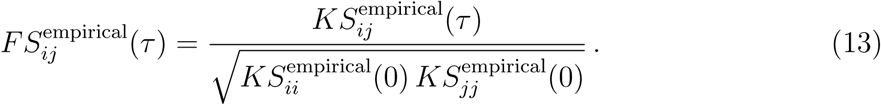

Note that these normalized time-shifted covariances break the symmetry of the couplings, thereby improving the quality of the fit [55, 56]. Using a heuristic pseudo-gradient algorithm, we iteratively updated 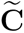 following the rule:

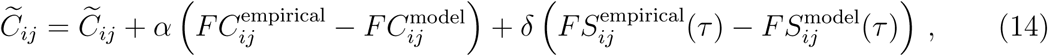

where 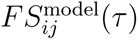 is defined similar to 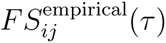. Details on the computation of the **KS** matrix can be found in [33]. We used *α* = *δ* = 1*e*^−5^ and continued the optimization until convergence. Importantly, updates to C were constrained by anatomical connectivity, meaning only the existing structural connections were allowed to take nonzero values. Details on the computation can be found in [34] and [33].

##### FDTest in a Hopf whole-brain model

We applied the FDTest to synthetic data generated with the Hopf whole-brain model. A step-like perturbation was applied simultaneously to all nodes in the network, and we then analyzed the local response of each oscillator separately. This protocol aims to mimic experimental neuroscience paradigms where a stimulus is applied to a wide area of the brain comprising multiple regions, while the response is recorded in various sites. The perturbation strength was set to *ε* = 0.02 and the perturbation was applied *M* = 5000 times. For each oscillator, we estimated the probability densities in the local 2D space, computed the conjugated variable, and calculated the mean transient response and the autocorrelation. Finally, we evaluated the local FDT deviation following Eq. 10.

##### Null models of brain connectivity

We investigated whether the FDT deviations in the brain are significantly different from what would be expected by chance in networks with shared features. To this end, we constructed three sets of null network models, each designed to preserve specific characteristics of the original brain connectivity: weight-distribution-preserving rewiring (RAND), strength-preserving rewiring (SPR), and topology-preserving rewiring (TPR). In the following, we provide a detailed description of each null model. The RAND model was obtained by a random rewiring of the original network, representing the least conservative model. The SPR was generated using the *netneurotools* Python library [57], which provides the function *in strength preserving rand sa dir* for generating surrogate networks that preserve both in-and out-strength sequences. Finally, the TPR was generated by random permutation of the weights without moving the edges. We generated 50 networks from each model and performed the FDTest as described in the previous section. Finally, we calculated the global FDT deviation of each network by averaging Δ^2^ across all nodes.

##### Linear stability and stationary network’s statistics

The statistics of the Hopf whole-brain model can be approximated by linearizing the dynamics around the origin, which is the fixed point [21]. The evolution of linear fluctuations *δ***u** = (*δ***x***, δ***y**) around the fixed point follows the stochastic linear equation:

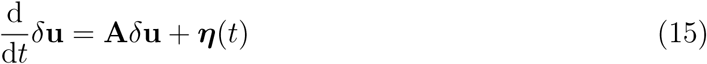

**A** is the Jacobian matrix at the fixed point, which can be written as a block matrix:

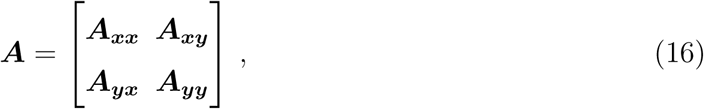

where ***A_xx_***= ***A_yy_***= diag(***a***−***S***)+***C*** and ***A_xy_***= −***A_yx_*** = diag(***ω***). Here ***a*** is the bifurcation parameter, ***S*** is the vector of node strengths *S_i_* = Ʃ*_j_ C_ij_* and ***ω*** is the vector of node intrinsic frequencies. Note that the Jacobian depends on the fitted effective connectivity.

To characterize the stability, we calculated the eigenvalues of the Jacobian matrix *λ_i_* for each fitted model. The origin is asymptotically stable if max [Re(*λ_i_*)] *<* 0, i.e. the largest real part of the set of eigenvalues of ***A*** is negative. In this work, all the fitted models were stable. However, in some models the value of max [Re(*λ_i_*)] closer to zero than others, meaning that the system was in the proximity of losing stability.

Additionally, we used the linear approximation to characterize the decay of the autocorrelation function of each node. The stationary covariance matrix of the fluctuations around the origin, i.e. ***C_v_***= ⟨*δ****u****δ****u****^T^* ⟩, of the linearized system (Eq. 15) is given by the Lyapunov equation:

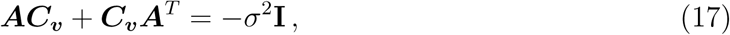

where the right-hand side is the covariance matrix of the uncorrelated, homogeneous noise. The Lyapunov equation can be solved numerically to obtain ***C_v_***. Furthermore, the stationary lagged covariances of the state variables, i.e. ***C_v_***(*τ*) = ⟨*δ****u***(*t* + *τ*)*δ****u****^T^* (*t*)⟩, can be obtained by:

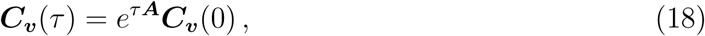

where ***C_v_***(0) = ***C_v_*** is the covariance matrix (zero-lag). The autocorrelation function (ACF) is the normalized lagged-autocovariance i.e. the diagonal elements of ***C_v_***(*τ*) normalized by ***C_v_***(0). The ACF is an exponentially decaying oscillating function. We characterized the speed of the decay by fitting the envelope of the ACF with an exponentially decaying function env|*C_v_*(*τ*)| ∼ *e*^−^*^τ/ξ^*. For each individualized model, we estimated the correlation length *ξ* of each node.

## ACKNOWLEDGMENTS

We thank Benjamin Lindner for the initial discussions on the generalized fluctuation-dissipation relation, which were instrumental in the development of this work. **S.M.G.** acknowledges support of AGAUR, Generalitat de Catalunya and Fondo Social Europeo (2022 FI B 00511). **J.M.M.**acknowledges financial support from CONICET (Argentina). **A.P.A.** is supported by a Ramón y Cajal fellowship (RYC2020-029117-I) funded by MI-CIU/AEI/10.13039/501100011033 and ESF Investing in your future, and by the Spanish State Research Agency, through the Severo Ochoa and Maŕıa de Maeztu Program for Cen-ters and Units of Excellence in R&D (CEX2020-001084-M). **Y.S.P.** was supported by the project NEurological MEchanismS of Injury, and Sleep-like cellular dynamics (NEMESIS) (ref. 101071900) funded by the EU ERC Synergy Horizon Europe. **G.D.** is supported by grant PID2022-136216NB-I00 funded by MICIU/AEI/10.13039/ 501100011033 and by “ERDF A way of making Europe,” ERDF, EU; Project Neurological Mechanisms of Injury and Sleep-like Cellular Dynamics (NEMESIS) (ref. 101071900), funded by the EU ERC Synergy Horizon Europe; and AGAUR research support grant (ref. 2021 SGR 00917), funded by the Department of Research and Universities of the Generalitat of Catalunya.

**S.M.G.**, **J.M.M.**, **A.P.A.**, **G.D.**, and **Y.S.P.** designed the research. **S.M.G.** and **J.M.M.** conducted the research. **S.M.G.**, **J.M.M.**, **A.P.A.**, **G.D.**, and **Y.S.P.** analyzed and interpreted the results. **Y.S.P.** and **G.D.** supervised the research. **S.M.G.** and **J.M.M** wrote the manuscript. All the authors edited the manuscript.

## DATA AVAILABILITY

Code to reproduce this paper has been deposited at zenodo and is openly available [58]

## APPENDIX

### DISAMBIGUATING MECHANISMS UNDERLYING FDT VIOLATIONS

In Figure 3 we showed that higher local FDT violations are associated with stronger connectivity. Here, we hypothesize that part of this effect is explained by the fact that the network drives the node’s effective dynamics far from the bifurcation, towards the noisy regime. Formally, from Eqs. 11 and 12 we can rewrite the model as:

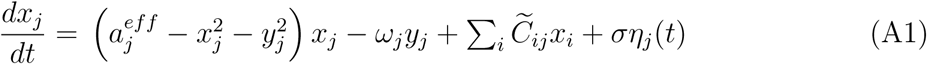

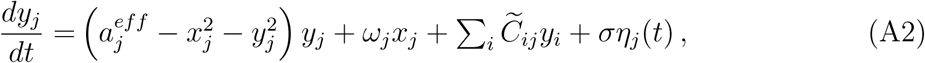

where 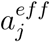 is the node’s effective bifurcation parameter

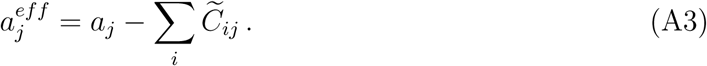

We then asked whether an isolated oscillator with an equivalent bifurcation parameter would exhibit the same FDT deviation. For each node of the network, we simulated a disconnected oscillator (Eqs. 7 and 8) using the node’s bifurcation parameter 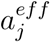 and frequency *w_j_*. The result, displayed in Figure 6A, indicates that a more negative bifurcation parameter (and thus higher strength) yields higher Δ^2^, in both connected and disconnected oscillators.

This result highlights the fact that Δ^2^ not only captures genuine FDT-violations, but also finite-sampling and histogram-estimation errors—that also depend on *a*. However, this effect is not sufficient to explain FDT deviations in the network, which resulted consistently greater. Specifically, nodes with lower connectivity strength —whose effective dynamics are near the bifurcation— exhibit deviations closer to their disconnected counterparts, while nodes with higher connectivity deviate more from the behavior predicted by their disconnected counterparts Fig. 6B. We find that network dynamics are necessary to explain the non-Markovian effects and their association with the connectivity strength.

**FIG. 6.**
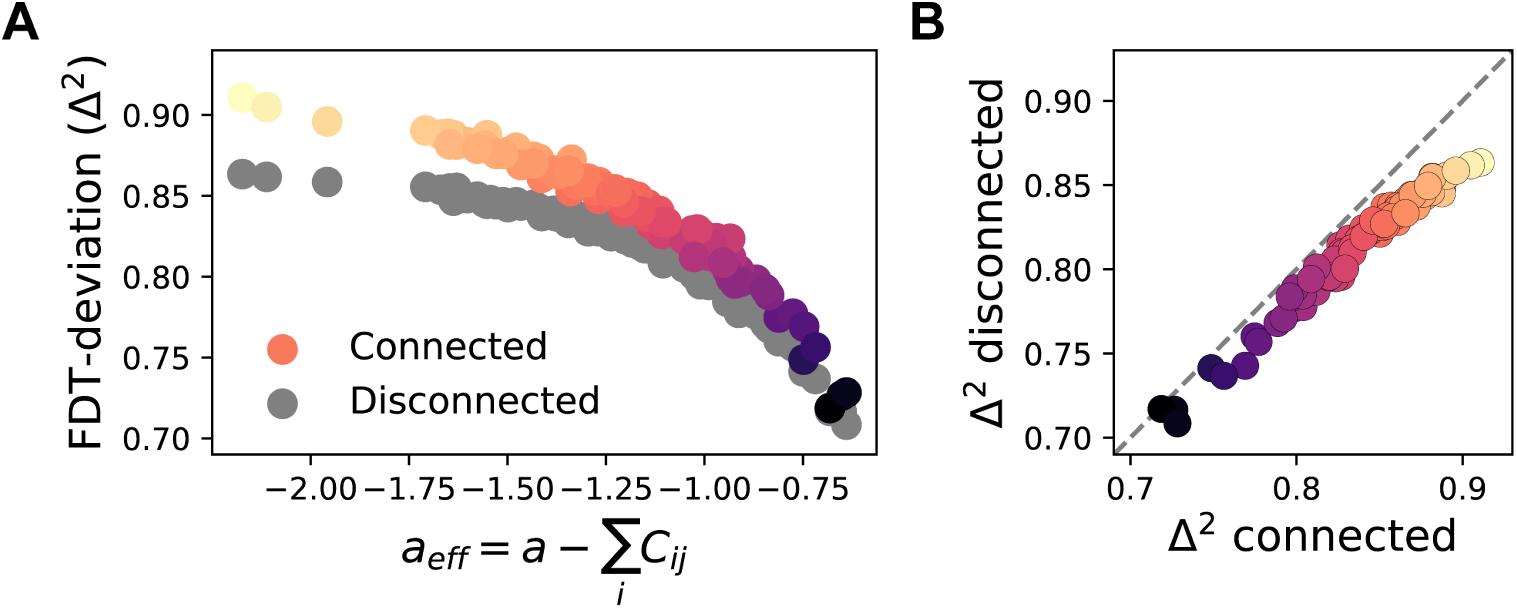
The effect of local dynamics in FDT deviations. **A**. FDT deviations as a function of the effective bifurcation parameter, for connected and equivalent disconnected oscillators. **B** FDT deviations in connected vs disconnected settings. Connected nodes show systematically higher deviations, especially those with strong coupling.

